# Global sensitivity analysis for investigation of bacterial physiology in complex media with FBA-PRCC

**DOI:** 10.1101/2024.02.14.580207

**Authors:** Anatoly Sorokin, Igor Goryanin

## Abstract

In this paper we propose the use of FBA-PRCC approach, which is based on sampling flux space and global sensitivity analysis (GSA), for analysis of the GEM FBA solution space in the nutrient-rich environment. We identify two new modes of interaction between species and metabolites: attraction and avoidance and propose the way of their use. We have shown that sensitivity coefficients provide additional information about behavior of the bacteria in the nutrient-rich environment in comparison to standard knockouts, FVA and CoPE-FBA analysis.

## Introduction

Flux-based analysis (FBA) of whole-genome metabolic reconstructions (GEM) stands as a computational method at the forefront of systems biology and metabolic engineering (Orth, Thiele, and Palsson 2010; Norsigian et al. 2019). This sophisticated approach plays a pivotal role in modeling and analyzing the intricate networks of biochemical reactions that constitute an organism’s metabolism. The essence of FBA lies in its ability to provide a quantitative framework, offering predictions and insights into the dynamic flow of metabolites through cellular metabolic pathways under specific environmental or physiological conditions (Thiele and Palsson 2010).

Whole-genome metabolic reconstructions (GEMs) serve as foundational datasets representing the entirety of biochemical reactions that an organism’s genome encodes. These reconstructions encompass a detailed inventory of metabolic pathways, including reactions, enzymes, and associated metabolites. GEMs are constructed based on the genomic information of an organism, and they serve as a comprehensive roadmap of its metabolic potential. Recent advances in the GEM reconstruction reviewed in (Fang, Lloyd, and Palsson 2020)

Flux-based analysis, when applied to GEMs, takes advantage of the stoichiometry and connectivity information embedded within these reconstructions. It operates on the principle of mass balance, considering the conservation of metabolites throughout the network. FBA enables the determination of flux distributions, representing the rates at which metabolites converts by different reactions within the cellular metabolism. The process begins by defining an objective function, typically the optimization of a specific cellular process, such as biomass production, ATP generation, or substrate utilization. FBA then employs linear programming techniques to find the optimal distribution of flux values through the metabolic network that maximizes or minimizes the chosen objective function, subject to the constraints imposed by the system (Orth, Thiele, and Palsson 2010).

In practical terms, FBA allows researchers to simulate and predict how an organism’s metabolism responds to various conditions. For example, FBA can be used to analyze how changes in nutrient availability, genetic modifications, or environmental factors influence the distribution of metabolic fluxes. This predictive capability is particularly valuable in metabolic engineering, where researchers aim to design or optimize microbial strains for the production of specific compounds, such as biofuels, pharmaceuticals, or industrial chemicals (Tian et al. 2017; Burgard, Pharkya, and Maranas 2003; McAnulty et al. 2012; Oyetunde et al. 2018; Sen 2022).

Another area of the active use of FBA models is in the medical application. Development of the human metabolic reconstruction (Ma et al. 2007; Thiele et al. 2013) and reorganization it with tissue specific submodels into integrated whole-body model (Thiele et al. 2020) open the avenue for a better understanding of the pathological processes such as cancer development (Gatto, Ferreira, and Nielsen 2020; Özcan and ÇakIr 2016). Apart from human GEM number of large collection of models were created by manual curation (Norsigian et al. 2019), semi-automated reconstruction (Magnúsdóttir et al. 2017; Heinken et al. 2023), or fully-automated reconstruction (Lieven et al. 2020) techniques.

What is common between all such reconstructions, as was noted by Van Pelt-KleinJan with coauthors (Van Pelt-KleinJan, De Groot, and Teusink 2021), is that all of them are based on experimental flux data from limited media growth. This is reasonable from experimental point of view, as single carbon source experiment allow to compare experimental growth rate with essentiality data from the model. But that type of model identification impose some limitations to the interpretation of the model results in nutrient-rich set up. The rich environment is very common as in biotechnological application, for example in the food industry (Mendoza Farías 2023; Qiu et al. 2023) and when bacteria participate in the community and interact with its members by production and consumption of nutrients (Kost et al. 2023). The interaction between pathways utilizing different substrates make solution space more complex, and increase number of non-unique solutions to the optimization FBA problem. In (Van Pelt-KleinJan, De Groot, and Teusink 2021) authors propose the enumeration of Elementary Conversion Modes technique to overcome that limitation. In this paper we propose the use of alternative approach (Sorokin and Goryanin 2023), which is based on sampling flux space and global sensitivity analysis (GSA), for analysis of the GEM FBA solution space in the nutrient-rich environment.

The randomized sampling of the FBA solution space is widely used for analysis of (Herrmann et al. 2019; Schellenberger and Palsson 2009). The great advantage of sampling is that it does not require any objective function to be optimized, so it allows to identify hidden relationships between reactions, genes and metabolites imposed by the physical, chemical and kinetic constraints applied to the metabolic network. That approach allow identification of the high-rate backbone of the model (Almaas et al. 2004), analysis of flux correlations and identification of flux modules (Papin, Reed, and Palsson 2004). Number of algorithms were developed for the sampling solution space (Fallahi, Skaug, and Alendal 2020). However, there are approaches of sampling, which are different from straightforward Monte-Carlo. Loghmani et al. (Loghmani et al. 2022) combine random flux sampling with Flux Variability Analysis (FVA) to separate reactions into “robust” and “sensitive” groups. In our FBA-PRCC approach (Sorokin and Goryanin 2023) we are sampling not the flux values space, but the parameter space of the FBA model which describes the reaction boundaries to identify boundaries that significantly influence the cost function via GSA techniques.

The Global Sensitivity Analysis (GSA) is an approach used in dynamic modelling to evaluate the impact of parameters on the model output (Marino et al. 2008; Lebedeva et al. 2012). Application of GSA techniques to the FBA GEM models recently gain interest, for example Damiani and co-authors implement Sobol’s variance based indices to understand how variation of exchange reaction constraint influences the growth rate (Damiani, Pescini, and Nobile 2020; Nobile et al. 2021).

In FBA-PRCC we apply Partial Rank Correlation GSA technique (Marino et al. 2008) in a similar way to the FBA model by vary flux boundaries and estimate its influence on target function. We called reaction sensitive if at least one of its boundary value show statistically significant effect on the target function. Unlike Damiani we optimized lysine production and estimate sensitivity coefficients of all reaction constraint to identify distant reactions influencing amino acid biosynthesis.

There are two key differences between Sobol’s variance based indices and Partial Rank Correlation coefficienst: (i) Sobol’s method is extremely computationally expensive, and (ii) it provides only value of the index, while PRCC gives both value and sign of the influence. Later, we will discuss how sensitivity coefficient sign could be interpret in terms of specie-metabolite interaction.

In this paper we will use growth rate as the FBA cost function and restrict ourselves to the analysis of media components, i.e. we will estimate sensitivity indices for the exchange reactions only.

## Results

For the analysis of the role of nutrient availability on the bacterial growth FBA uses so-called extracellular metabolites and exchange reactions. Flux through exchange reaction represents consumption of the compound if its value is negative and production if it is positive. So, for all metabolites that could be consumed by the cell exchange reactions have negative lower bound and for all metabolites that could be exported from the cell corresponding exchange reaction have positive upper boundary. To visualize this, we could draw arrow from cell to metabolite for production flux and from metabolite to cell for consumption flux (Fig. 1). We sampled absolute value of each non-zero constraints of exchange reactions and analyzed nutrient sensitivity of the Escherichia coli K12 sub-strain MG1655 GEM from AGORA2 collection (Heinken et al. 2023) and two autotrophic mutants ΔhisD and ΔtrpB, which are deficient in biosynthesis of histidine and tryptophan correspondently. For each PRC coefficient we have estimate its significance and kept only those with p-value below 0.1. Out of 1114 constraints in the model only 102 are significant at this level out of which 49 exchange reactions have lower bound significant (substrates) and 53 have upper bound significant (products). Four metabolites have both constraints significantly sensitive: Acetaldehyde, Cholate, Ethanol, and Trimethylamine, which means that production and consumption significantly influences growth rate. Majority of the exchange reactions (89) have positive sensitivity coefficients and only 13 have negative one (Table 1).

**Table 1.** Sign distribution among sensitivity coefficients

| Sign/Boundary | Lower | Upper |
| --- | --- | --- |
| Negative | 11 | 2 |
| Positive | 38 | 51 |

**Figure 1.**
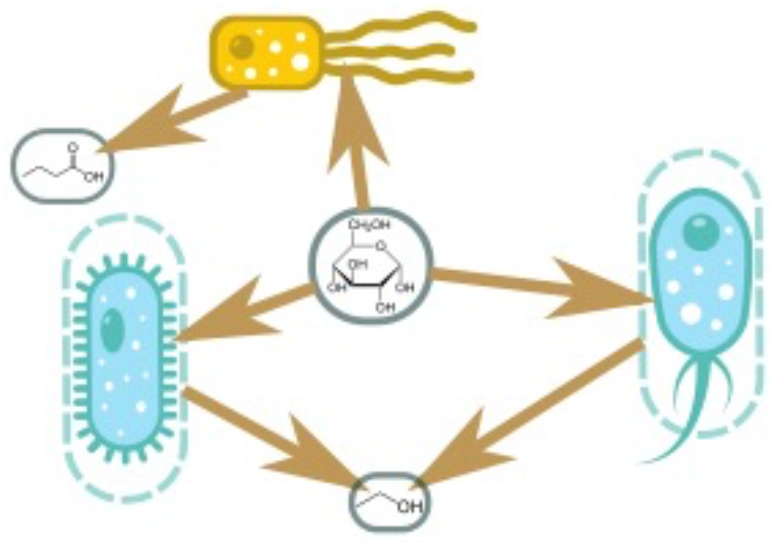
Schematic representation of the specie-metabolite network.

The exchange reaction in the standard FBA paradigm represents passive diffusion so interpretation of the positive PRCC values is straightforward. Positive sensitivity of the upper bound indicate that the growth positively correlates with the rate of metabolite production, so we could call such metabolite product. In a similar manner positive sensitivity coefficient of lower bound means that growth significantly depends upon consumption of the corresponding metabolite (Fig.2A).

Interpretation of the negative coefficient is more challenging. Negative concentration for the upper/lower bound indicate that increase in rate of the metabolite diffusion from/to the cell corresponds to decrease of the growth. In that situation cell would try to decrease the corresponding diffusion rate, and as the diffusion rate is defined by the concentration gradient the obvious way to decrease the rate is to decrease the gradient. For the upper bound it would mean to find the niche with high concentration of the corresponding metabolite, so cell could still produce it but won’t lose it via diffusion. For the lower bound it would mean to find the niche with low concentration of the correspondent metabolite, so there is nothing to consume. According to this line of reason we will call metabolites with negative upper bound PRCC value attractant and metabolites with negative lower bound PRCC – repellent (Fig. 2B).

**Figure 2.**
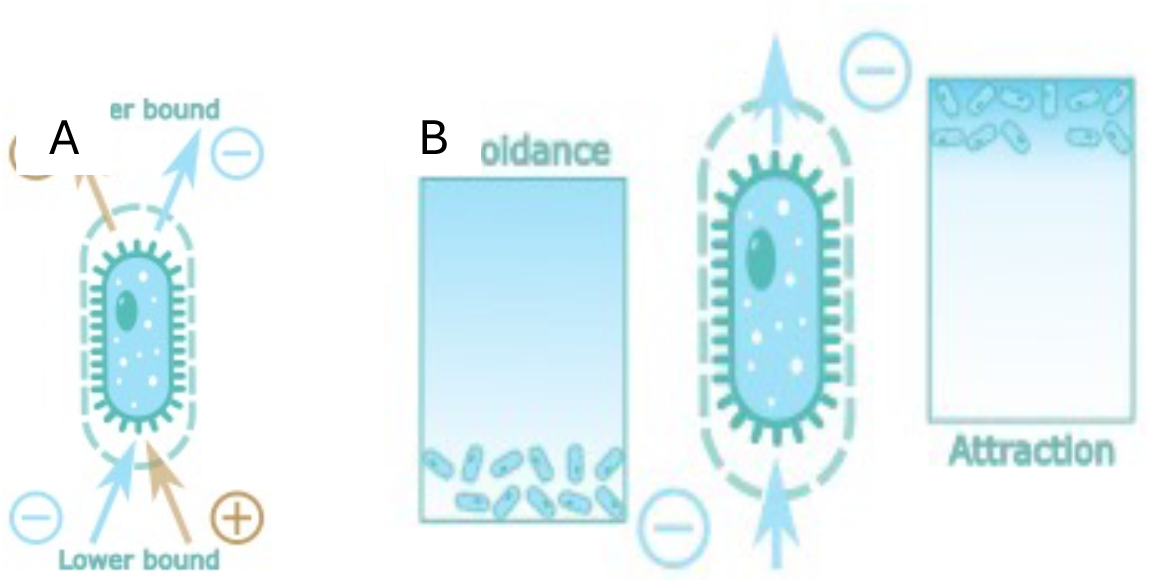
FBA-PRCC sign interpretation.

Table 1 shows that majority of the metabolites have positive PRCC coefficients and represent standard types of specie-metabolite interactions parts. There are only two attractants (M3 Glucuronide and regorafenib-N-ß-glucuronide) both of which are exometabolites from drug metabolic pathways. Their sensitivity most probably due to incomplete metabolization pathway rather than real effect on the cell. There are 11 repellents (Table 2). Out of them two metabolites Cholate and Ethanol have negative lower boundary and positive upper one, which means that they are toxic byproducts which good to be produced but not so good to have around. Another two metabolites have significant positive sensitivity coefficients on both boundaries values: Acetaldehyde and Trimethylamine. That probably indicate presence of the futile transport cycle in the model, when important metabolite excreted and then imported again.

**Table 2.**
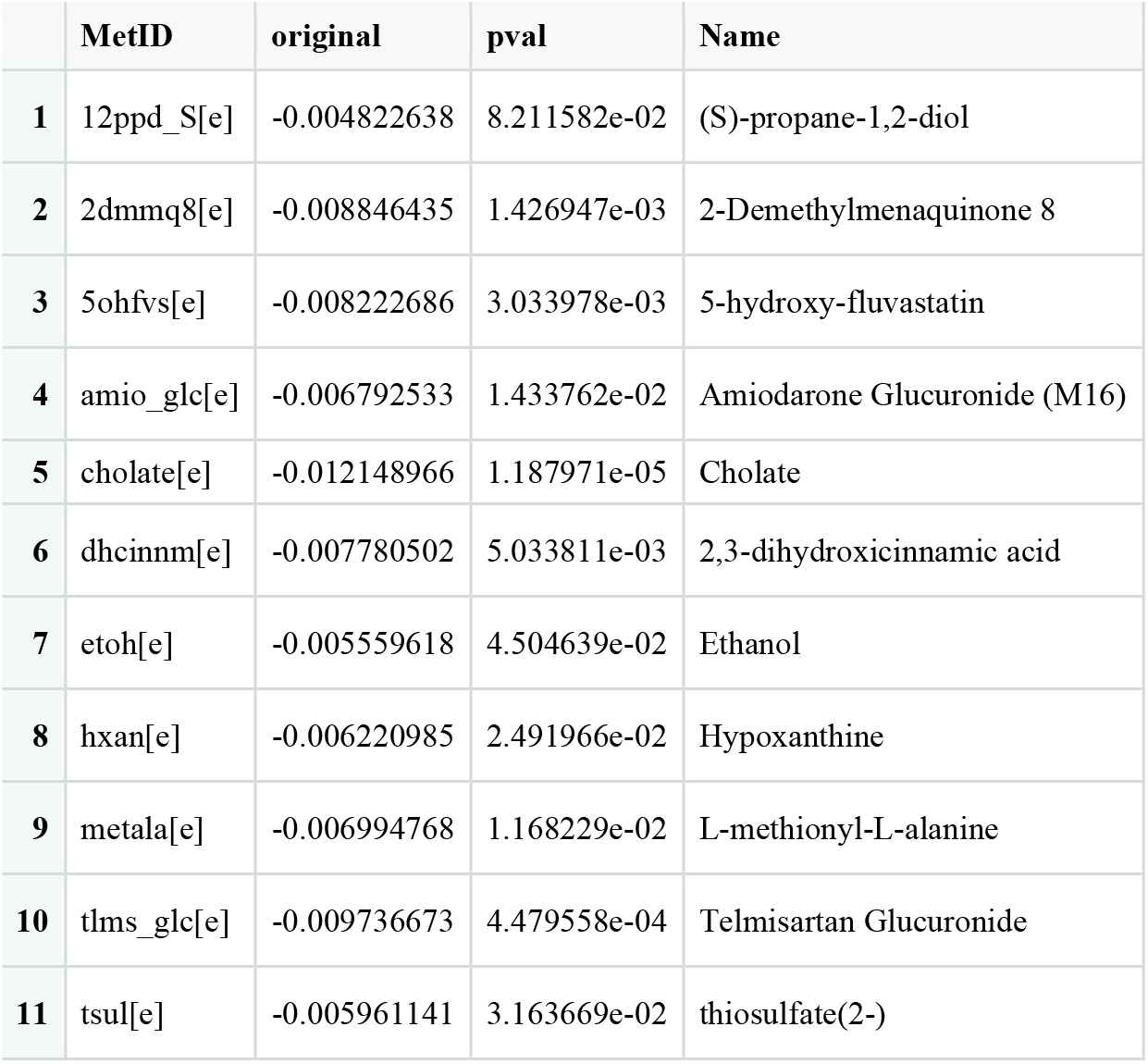

## Materials and Methods

The metabolic network with *m* metabolites and *r* reactions is described by a *m · r* stoichiometry matrix *N*. The (*i, j*)-th entry of *N*, n_ij_, is the stoichiometric coefficient of the *i*-th metabolite in the *j*-th reaction. Any reaction flux vector *v* that satisfies *Nv*=0 contains reaction fluxes such that the system is in a steady state. In Flux Balance Analysis (FBA) [18] some optimization problem is solved to identify unique solution vector v_o_ such that 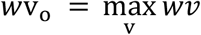 for v_1_≤ v ≤ v_u_, where *w* is the objective coefficient vector, and reaction bounds are V_1_ and v_u_. We are interested in the estimation of the sensitivity of the objective function to the values of reaction boundaries.

There is a special type of reaction in the constraint-based modelling called ‘boundary reactions’, which usually describe the exchange of metabolites between the internal ‘cell’ and the external ‘environment’.

Our approach consists of three steps:

1. Define parameter space: for irreversible boundary reactions only one parameter v_u_ is created, for reversible boundary reactions two parameters created for each reaction – v_1_ and v_u_.
2. Generate a set of quasi-random low-discrepancy points in the parameter space. Update parameters (reaction bounds) and find the optimal objective value for each point in the parameter space.
3. Calculate Partial Rank Correlation Coefficient (PRCC) for each parameter and objective values. The statistical significance of the PRCC value is estimated by as described by Marino et al (Marino et al. 2008). The sufficiency of the sample size for reliable PRCC estimation is controlled by the top-down coefficient of concordance (TDCC): when TDCC between PRCC vectors calculated at different sample sizes exceed the threshold of 0.9, sample size is considered sufficient for analysis.

The model E. coli str. K-12 substr. MG1655 was taken from the AGORA2 collection (Heinken et al. 2023) in SBML format. Mutant model ΔhisD was created by removing reactions HISTLOX and HISTDOX which controlled by hisD gene, encoding Histidinol dehydrogenase. Mutant model ΔtrpB was created by removing reactions TRPS1, TRPS2, and TRPS3, which depends upon trpB gene encoding tryptophan synthase α-chain.

The random sampling of the exchange model constraint was performed with the Sobol quasi-random low-discrepancy sequence (Renardy et al. 2021; Marino et al. 2008) that was generated with the python Quasi-Monte Carlo submodule of the SciPy v.1.7.3 (Virtanen et al. 2020). Model simulation was performed with the Cobrapy v.0.25.0 (Ebrahim et al. 2013). All calculations are performed with Python version 3.10.2.

PRCC calculations were performed with R package ‘sensitivity’ version 1.28.0 (Iooss et al. 2022). TDCC values are calculated by ‘ODEsensitivity’ R package version 1.1.2 (Weber and Theers 2019). All calculations are performed with R version 4.2.1 (R Core Team 2022).

All simulations were performed on the OIST HPC cluster with 8CPU and 64GB per job. Sobol points generation, application to the reaction boundaries and optimization of objectives were performed in chunks of 8192 per job. Calculations of the PRCC sensitivity coefficients were performed on 262144 Sobol points in chunks of 10 features per job. Convergence of the calculation was controlled by TDCC between consecutive datasets different in 8192 Sobol points. The TDCC value between 262144 and 253952 was 0.909. The average execution time is 30 minutes per job for the Sobol point calculations and 7 hours per job for the PRCC calculation.

## Discussion

Mechanistic understanding of role of different components of the rich media for the growth and physiology of microorganisms is of paramount importance for solving both scientific and technological problems. Overflow metabolism, which cause overproduction of byproducts in the nutrient-rich environment, could decrease yield in biotechnological reactor and make specie keystone in the complex microbial community. In this paper we have shown that two new modes of specie-metabolite interaction could be added to the standard consumption and production. Avoidance interaction describes the interaction in which cell looking for the environment with low concentration of the metabolite, for example, some microaerophile strains demonstrate avoidance link with the oxygen (data not shown). Attraction interaction on the other side makes cell look for the environment with high concentration of the metabolite, for example to prevent lost of the important intermediate via passive transport.

Proposed analysis opens a new way of looking into the specie-metabolite interactions and their role in microbial community dynamics and single specie physiology.

## Acknowledgments

We are grateful for the help and support provided by the Scientific Computing and Data Analysis section of Research Support Division at OIST.

## Reference

Almaas, E., B. Kovács, T. Vicsek, Z. N. Oltvai, and A.-L. Barabási. 2004. ‘Global Organization of Metabolic Fluxes in the Bacterium Escherichia Coli’. Nature 427 (6977): 839–43. 10.1038/nature02289.

Burgard, Anthony P., Priti Pharkya, and Costas D. Maranas. 2003. ‘Optknock: A Bilevel Programming Framework for Identifying Gene Knockout Strategies for Microbial Strain Optimization’. Biotechnology and Bioengineering 84 (6): 647–57. 10.1002/bit.10803.

Damiani, Chiara, Dario Pescini, and Marco S. Nobile. 2020. ‘Global Sensitivity Analysis of Constraint-Based Metabolic Models’. In Computational Intelligence Methods for Bioinformatics and Biostatistics, edited by Maria Raposo, Paulo Ribeiro, Susana Sério, Antonino Staiano, and Angelo Ciaramella, 11925:179–86. Lecture Notes in Computer Science. Cham: Springer International Publishing. 10.1007/978-3-030-34585-3_16.

Ebrahim, Ali, Joshua A Lerman, Bernhard O Palsson, and Daniel R Hyduke. 2013. ‘COBRApy: COnstraints-Based Reconstruction and Analysis for Python’. BMC Systems Biology 7 (1): 74. 10.1186/1752-0509-7-74.

Fallahi, Shirin, Hans J. Skaug, and Guttorm Alendal. 2020. ‘A Comparison of Monte Carlo Sampling Methods for Metabolic Network Models’. Edited by Inés P. Mariño. PLOS ONE 15 (7): e0235393. 10.1371/journal.pone.0235393.

Fang, Xin, Colton J. Lloyd, and Bernhard O. Palsson. 2020. ‘Reconstructing Organisms in Silico: Genome-Scale Models and Their Emerging Applications’. Nature Reviews Microbiology 18 (12): 731–43. 10.1038/s41579-020-00440-4.

Gatto, Francesco, Raphael Ferreira, and Jens Nielsen. 2020. ‘Pan-Cancer Analysis of the Metabolic Reaction Network’. Metabolic Engineering 57 (January): 51–62. 10.1016/j.ymben.2019.09.006.

Heinken, Almut, Johannes Hertel, Geeta Acharya, Dmitry A. Ravcheev, Malgorzata Nyga, Onyedika Emmanuel Okpala, Marcus Hogan, et al. 2023. ‘Genome-Scale Metabolic Reconstruction of 7,302 Human Microorganisms for Personalized Medicine’. Nature Biotechnology, January. 10.1038/s41587-022-01628-0.

Herrmann, Helena A., Beth C. Dyson, Lucy Vass, Giles N. Johnson, and Jean-Marc Schwartz. 2019. ‘Flux Sampling Is a Powerful Tool to Study Metabolism under Changing Environmental Conditions’. Npj Systems Biology and Applications 5 (1): 32. 10.1038/s41540-019-0109-0.

Iooss, Bertrand, Sebastien Da Veiga, Alexandre Janon, Gilles Pujol, with contributions from Baptiste Broto, Khalid Boumhaout, Thibault Delage, et al. 2022. ‘Sensitivity: Global Sensitivity Analysis of Model Outputs’. Manual. https://CRAN.R-project.org/package=sensitivity.

Kost, Christian, Kiran Raosaheb Patil, Jonathan Friedman, Sarahi L. Garcia, and Markus Ralser. 2023. ‘Metabolic Exchanges Are Ubiquitous in Natural Microbial Communities’. Nature Microbiology, November. 10.1038/s41564-023-01511-x.

Lebedeva, Galina, Anatoly Sorokin, Dana Faratian, Peter Mullen, Alexey Goltsov, Simon P. Langdon, David J. Harrison, and Igor Goryanin. 2012. ‘Model-Based Global Sensitivity Analysis as Applied to Identification of Anti-Cancer Drug Targets and Biomarkers of Drug Resistance in the ErbB2/3 Network’. European Journal of Pharmaceutical Sciences 46 (4): 244–58. 10.1016/j.ejps.2011.10.026.

Lieven, Christian, Moritz E. Beber, Brett G. Olivier, Frank T. Bergmann, Meric Ataman, Parizad Babaei, Jennifer A. Bartell, et al. 2020. ‘MEMOTE for Standardized Genome-Scale Metabolic Model Testing’. Nature Biotechnology 38 (3): 272–76. 10.1038/s41587-020-0446-y.

Loghmani, Seyed Babak, Nadine Veith, Sven Sahle, Frank T. Bergmann, Brett G. Olivier, and Ursula Kummer. 2022. ‘Inspecting the Solution Space of Genome-Scale Metabolic Models’. Metabolites 12 (1): 43. 10.3390/metabo12010043.

Ma, Hongwu, Anatoly Sorokin, Alexander Mazein, Alex Selkov, Evgeni Selkov, Oleg Demin, and Igor Goryanin. 2007. ‘The Edinburgh Human Metabolic Network Reconstruction and Its Functional Analysis’. Molecular Systems Biology 3 (1): 135. 10.1038/msb4100177.

Magnúsdóttir, Stefanía, Almut Heinken, Laura Kutt, Dmitry A Ravcheev, Eugen Bauer, Alberto Noronha, Kacy Greenhalgh, et al. 2017. ‘Generation of Genome-Scale Metabolic Reconstructions for 773 Members of the Human Gut Microbiota’. Nature Biotechnology 35 (1): 81–89. 10.1038/nbt.3703.

Marino, Simeone, Ian B. Hogue, Christian J. Ray, and Denise E. Kirschner. 2008. ‘A Methodology for Performing Global Uncertainty and Sensitivity Analysis in Systems Biology’. Journal of Theoretical Biology 254 (1): 178–96. 10.1016/j.jtbi.2008.04.011.

McAnulty, Michael J, Jiun Y Yen, Benjamin G Freedman, and Ryan S Senger. 2012. ‘Genome-Scale Modeling Using Flux Ratio Constraints to Enable Metabolic Engineering of Clostridial Metabolism in Silico’. BMC Systems Biology 6 (1): 42. 10.1186/1752-0509-6-42.

Mendoza Farías, Sebastián Nelson. 2023. ‘Metabolic Modeling of Microorganisms and Dynamics of Microbial Communities in the Food Industry’. PhD, Vrije Universiteit Amsterdam. 10.5463/thesis.540.

Nobile, Marco S., Vasco Coelho, Dario Pescini, and Chiara Damiani. 2021. ‘Accelerated Global Sensitivity Analysis of Genome-Wide Constraint-Based Metabolic Models’. BMC Bioinformatics 22 (S2): 78. 10.1186/s12859-021-04002-0.

Norsigian, Charles J, Neha Pusarla, John Luke McConn, James T Yurkovich, Andreas Dräger, Bernhard O Palsson, and Zachary King. 2019. ‘BiGG Models 2020: Multi-Strain Genome-Scale Models and Expansion across the Phylogenetic Tree’. Nucleic Acids Research, November, gkz1054. 10.1093/nar/gkz1054.

Orth, Jeffrey D, Ines Thiele, and Bernhard Ø Palsson. 2010. ‘What Is Flux Balance Analysis?’ Nature Biotechnology 28 (3): 245–48. 10.1038/nbt.1614.

Oyetunde, Tolutola, Forrest Sheng Bao, Jiung-Wen Chen, Hector Garcia Martin, and Yinjie J. Tang. 2018. ‘Leveraging Knowledge Engineering and Machine Learning for Microbial Bio-Manufacturing’. Biotechnology Advances 36 (4): 1308–15. 10.1016/j.biotechadv.2018.04.008.

Özcan, Emrah, and Tunahan ÇakIr. 2016. ‘Reconstructed Metabolic Network Models Predict Flux-Level Metabolic Reprogramming in Glioblastoma’. Frontiers in Neuroscience 10 (April). 10.3389/fnins.2016.00156.

Papin, J, J Reed, and B Palsson. 2004. ‘Hierarchical Thinking in Network Biology: The Unbiased Modularization of Biochemical Networks’. Trends in Biochemical Sciences 29 (12): 641–47. 10.1016/j.tibs.2004.10.001.

Qiu, Sizhe, Hong Zeng, Zhijie Yang, Wei-Lian Hung, Bei Wang, and Aidong Yang. 2023. ‘Dynamic Metagenome-scale Metabolic Modeling of a Yogurt Bacterial Community’. Biotechnology and Bioengineering 120 (8): 2186–98. 10.1002/bit.28492.

R Core Team. 2022. ‘R: A Language and Environment for Statistical Computing’. Manual. Vienna, Austria. https://www.R-project.org/.

Renardy, Marissa, Louis R. Joslyn, Jess A. Millar, and Denise E. Kirschner. 2021. ‘To Sobol or Not to Sobol? The Effects of Sampling Schemes in Systems Biology Applications’. Mathematical Biosciences 337 (July): 108593. 10.1016/j.mbs.2021.108593.

Schellenberger, Jan, and Bernhard Ø. Palsson. 2009. ‘Use of Randomized Sampling for Analysis of Metabolic Networks’. Journal of Biological Chemistry 284 (9): 5457–61. 10.1074/jbc.R800048200.

Sen, Pramita. 2022. ‘Flux Balance Analysis of Metabolic Networks for Efficient Engineering of Microbial Cell Factories’. Biotechnology and Genetic Engineering Reviews, December, 1–34. 10.1080/02648725.2022.2152631.

Sorokin, Anatoly, and Igor Goryanin. 2023. ‘FBA-PRCC. Partial Rank Correlation Coefficient (PRCC) Global Sensitivity Analysis (GSA) in Application to Constraint-Based Models’. Biomolecules 13 (3): 500. 10.3390/biom13030500.

Thiele, Ines, and Bernhard Ø Palsson. 2010. ‘A Protocol for Generating a High-Quality Genome-Scale Metabolic Reconstruction’. Nature Protocols 5 (1): 93–121. 10.1038/nprot.2009.203.

Thiele, Ines, Swagatika Sahoo, Almut Heinken, Johannes Hertel, Laurent Heirendt, Maike K Aurich, and Ronan MT Fleming. 2020. ‘Personalized Whole-body Models Integrate Metabolism, Physiology, and the Gut Microbiome’. Molecular Systems Biology 16 (5). 10.15252/msb.20198982.

Thiele, Ines, Neil Swainston, Ronan M T Fleming, Andreas Hoppe, Swagatika Sahoo, Maike K Aurich, Hulda Haraldsdottir, et al. 2013. ‘A Community-Driven Global Reconstruction of Human Metabolism’. Nature Biotechnology 31 (5): 419–25. 10.1038/nbt.2488.

Tian, Mingyuan, Prashant Kumar, Sanjan T. P. Gupta, and Jennifer L. Reed. 2017. ‘Metabolic Modeling for Design of Cell Factories’. In Systems Biology, edited by Jens Nielsen and Stefan Hohmann, 1st ed., 71–107. Weinheim, Germany: Wiley-VCH Verlag GmbH & Co. KGaA. 10.1002/9783527696130.ch3.

Van Pelt-KleinJan, Eunice, Daan H. De Groot, and Bas Teusink. 2021. ‘Understanding FBA Solutions under Multiple Nutrient Limitations’. Metabolites 11 (5): 257. 10.3390/metabo11050257.

Virtanen, Pauli, Ralf Gommers, Travis E. Oliphant, Matt Haberland, Tyler Reddy, David Cournapeau, Evgeni Burovski, et al. 2020. ‘SciPy 1.0: Fundamental Algorithms for Scientific Computing in Python’. Nature Methods 17 (3): 261–72. 10.1038/s41592-019-0686-2.

Weber, Frank, and Stefan Theers. 2019. ‘ODEsensitivity: Sensitivity Analysis of Ordinary Differential Equations’. Manual. https://CRAN.R-project.org/package=ODEsensitivity.

